# Invasive streptococcal infection can lead to the generation of cross-strain opsonic antibodies

**DOI:** 10.1101/2022.06.27.497877

**Authors:** Therese de Neergaard, Anna Bläckberg, Hanna Ivarsson, Sofia Thomasson, Vibha Kumra Ahnlide, Sounak Chowdhury, Hamed Khakzad, Johan Malmström, Magnus Rasmussen, Pontus Nordenfelt

## Abstract

**Introduction:** The human pathogen *Streptococcus pyogenes* causes substantial morbidity and mortality. It is unclear if antibodies developed after infections with this pathogen are opsonic and if they are strain-specific or more broadly protective. Here, we quantified the opsonic antibody response following invasive *S. pyogenes* infection.

**Materials and Methods:** Four patients with *S. pyogenes* bacteremia between 2018-2020 at Skåne University Hospital in Lund, Sweden, were prospectively enrolled. Acute and convalescent sera were obtained, and the *S. pyogenes* isolates were genome-sequenced (*emm*118, *emm*85, and two *emm*1). Quantitative antibody binding and phagocytosis assays were used to evaluate isolate-dependent opsonic antibody function in response to infection.

**Results:** Antibody binding increased modestly against the infecting isolate and across *emm* types in convalescent compared to acute sera for all patients. For two patients, phagocytosis increased in convalescent serum for both the infecting isolate and across types. The increase was only across types for one patient, and one had no improvement. No correlation to the clinical outcomes was observed.

**Conclusion:** Invasive *S. pyogenes* infections result in a modestly increased antibody binding with differential opsonic capacity, both non-functional binding and broadly opsonic binding across types. These findings question the dogma that an invasive infection should lead to a strong type-specific antibody increase rather than a more modest but broadly reactive response, as seen in these patients. Furthermore, our results indicate that an increase in antibody titers might not be indicative of an opsonic response and highlight the importance of evaluating antibody function in *S. pyogenes* infections.

## Introduction

*Streptococcus pyogenes,* Group A streptococcus (GAS), is each year estimated to cause more than 700 million mild skin and throat infections and around 600 000 invasive ones such as sepsis and necrotizing fasciitis (Carapetis et al., 2005). The substantial morbidity and mortality caused by *S. pyogenes* makes it important to understand the immune response to this pathogen.

A crucial part of the defense against pathogens is opsonizing antibodies, which, when bound to the pathogen, enhance its eradication by phagocytosis. However, *S. pyogenes* has evolved multiple strategies to resist phagocytosis (Carlsson et al., 2003; Fischetti, 1989; Staali et al., 2006). A major virulence factor in this process is the streptococcal M protein, encoded by the *emm* gene. M protein can reverse antibody orientation through Fc binding (Åkesson et al., 1994; Nordenfelt et al., 2012), interact with multiple anti-phagocytic proteins (Carlsson et al., 2005; Happonen et al., 2019), and exhibits antigenic diversity through its hypervariable region resulting in >250 *emm* types (Castro & Dorfmueller, 2021). The M protein covers most of the bacterial surface and is an important target for the immune system through type-specific antibodies. These antibodies start to appear around four weeks after a GAS infection (Denny et al., 1957) and persist up to 30 years (Bencivenga et al., 2009; Lancefield, 1959). It is generally believed that immunity is *emm* type-specific and is acquired through the development of protective type-specific antibodies. Initial studies suggested that only antibodies against the hypervariable part of the M protein were opsonic (Jones & Fischetti, 1988), but later studies have reported that antibodies to conserved binding sites can also be opsonic (Bahnan et al., 2021; Pandey et al., 2019; Vohra et al., 2005). These findings indicate the presence of anti-M antibodies, which may convey immunity to more than one *emm* type providing a broader protection. The immune response might also target other parts of the bacteria, such as carbohydrates of the cell wall (Gao et al., 2021), which could convey more general cross-type immunity. Early in life, children and adolescents suffer from recurring GAS infections; however, these infections decrease radically in adulthood. Therefore, it is suggested that through those repeated exposures over time, a more broad and long-term immunity is developed(Pandey et al., 2016).

Specific antibodies do not always activate immune functions, and antibody responses can be described as opsonic or non-opsonic. In contrast to its opsonic counterpart, a non-opsonic antibody binds to its antigen without contributing to eradicating the pathogen by phagocytosis (Bahnan et al., 2021; Bläckberg et al., 2021; Forthal, 2015). Recently, Bläckberg *et al.* reported that patients suffering from an invasive *Streptococcus dysgalactiae* infection failed to develop protective opsonic antibodies (Bläckberg et al., 2021), and Uddén *et al.* found that the generation of non-opsonic antibody responses was correlated to the invasive nature of the *Streptococcus pneumoniae* infection (Uddén et al., 2020). However, it is unknown to what extent this occurs for *S. pyogenes* infections and how important the nature of the infection, and in particular invasive disease, is in developing opsonizing antibodies to *S. pyogenes*.

To better understand the functional immune response in *S. pyogenes* infections, we assess antibody binding and their opsonic capacity in four patients during and after an invasive*S. pyogenes* infection. Interestingly, we report the development of both non-opsonic type-specific antibodies as well as broad opsonic antibody responses across different *emm*-types in these patients.

## Methods

### Patient inclusion and data collection

Patients with *S. pyogenes* bacteremia in 2018-2020 were prospectively included in the study after obtaining oral and written consent. Acute serum was collected within five days after hospital admission, and convalescent serum was collected after 4-6 weeks. Medical records of patients were reviewed to obtain clinical and epidemiological parameters. The concentration of the immunoglobulins in serum was determined at the Department of Clinical Chemistry in Skåne, Sweden.

### Ethics

The regional ethics committee approved the study of Lund University (2016/939, with amendment 2018/828).

### Sequencing

The *S. pyogenes* blood isolates were collected from the Laboratory for Clinical Microbiology, Lund University Hospital Sweden. Whole-genome sequencing was done at the Center for Translational Genomics at Lund University. NextSeq 550 Illumina sequencing was used to sequence the bacterial genomes. The genome sequencing data were searched against the CDC database of M protein families to detect the target M protein sequence. M protein sequences were pairwise aligned with the target M1 protein using EMBOSS Needle web server.

### Microbe strains

The clinical isolates of *Streptococcus pyogenes* and the lab mutant of *S. pyogenes* SF370 with deficient M protein expression (dM) (Abbot et al., 2007; Ferretti et al., 2001) were statically cultured in Todd Hewitt Broth (Bacto) supplemented with 0.2% yeast extract (Difco) (THY) at 37°C and 5% CO_2_. They were cultivated to log phase (OD_600 nm_ 0.3-0.4, Ultrospec 10; Amersham Biosciences) before being heat-killed at 80°C for 5 min.

### Labeling and Opsonization of bacteria

Bacteria in PBS were first stained for 1 h at 37°C with 4 μM Oregon Green 488-X succinimidyl ester (Invitrogen) followed by 20 μg/ml CypHer5E (Cytiva) in Na2CO3 for the phagocytosis assay. After staining the bacteria were resuspended in Na-medium (5.6 mM glucose, 127 mM NaCl, 10.8 mM KCl, 2.4 mM KH_2_PO_4_, 1.6 mM MgSO_4_, 10 mM HEPES, 1.8 mM CaCl_2_; pH adjusted to 7.3 with NaOH). To disperse any large aggregates, stained bacteria were then sonicated for 4 min (0.5 cycle, 75 A, VialTweeter) followed by determination of concentration using flow cytometry (CytoFlex; Beckman Coulter, lasers: 488 nm, 638 nm, filters: 525/40, 660/10)).

Opsonization was performed on the experiment day with a bacterial concentration of 800 000 bacteria/μl at 37°C for 30 min with gentle shaking. Sera were heat-inactivated before opsonization at 56°C for 30 min. The opsonin used besides patient sera were intravenous immunoglobulin pooled from healthy donors (IVIG; Octagam, Octapharma) and a humanized monoclonal IgG that is IgE-specific (Xolair, Omalizumab, Novartis) and thus only binds to M protein via potential Fc binding.

### Binding assay

Oregon Green stained bacteria were opsonized in a 1:2 serial dilution of sera, IVIG (2mg/ml), and Xolair (1mg/ml) in a final volume of 10 μl. For the assessment across the different isolates, bacteria were double-stained (Oregon Green, CypHer5E) and opsonized with 5% sera or 0.5 mg/ml of Xolair or IVIG. After opsonization, unbound antibodies were washed away three times by removing supernatant and resuspend in 250 μl Na-medium through centrifugation (3000 g, 5 min). Opsonized bacteria were stained with fluorescently labeled antibodies (Alexa Fluor 647-conjugated F(ab’)2 Fragment Goat Anti-Human IgG Fab; Jackson ImmunoResearch Laboratories) at 1:50 dilution for 30 min at 37°C. Data were obtained through CytoFlex, acquiring at least 15 000 events. Four separate bacterial colonies per isolate plate were picked and assessed.

### Affinity model

Binding curves were analyzed using a GAS antibody binding model based on the transfer matrix method for competitive binding described in Kumra Ahnlide et al., 2021. The implementation of this model is available on Github (10.5281/zenodo.4063760). Using this model, the binding of polyclonal antibody samples is characterized by the mean and range (95 % CI) of a log-normal distribution of affinities. The geometric means as given in the main figures correspond to the antibody affinity of the polyclonal samples. Binding values were normalized to an interpolated saturation level before being evaluated with the model implementation. Measured binding curves are shown as the mean and standard deviation of data points as described in the figure legends. Affinity values were derived by minimizing the weighted mean squared error of the model output and measured data using a MATLAB minimization function. The accuracy of predicted affinities was estimated using the bootstrap method, where the confidence intervals were calculated from 50 re-samplings of the measured data.

### Determination of immunoglobulin sub-classes

Mass spectrometry analysis was performed on patient sera to measure the immunoglobulin subclasses. The sample preparation for mass spectrometry is described elsewhere (Chowdhury et al., 2021). In brief, 8 M urea-100 mM ammonium bicarbonate was added to 1 μl of patient serum for denaturation, and 5 mM Tris(2-carboxyethyl) phosphine hydrochloride (TCEP) was added and incubated for 1 hour at 37°C for reduction, followed by incubation with 10 mM iodoacetamide for alkylation at room temperature for 30 minutes. Samples were diluted in 100 mM ammonium bicarbonate and incubated overnight with 0.5 μg/μl sequencing-grade trypsin (Promega) at 37°C, after which the addition of 10% formic acid inactivated trypsin. SOLAμ horseradish peroxidase (HRP) 2 mg/1 ml 96-well plate (Thermo Scientific) was used to concentrate the peptides (according to the manufacturer’s instructions). The concentrated peptides were injected into in a Q Exactive HFX instrument (Thermo-Scientific) connected to an Easy-nLC 1200 instrument (Thermo Scientific). The peptides were analyzed in data-dependent mass-spectrometry (DDA-MS) mode (Chowdhury et al., 2021). In short, the peptides were separated on a 50-cm Easy-Spray column (column temperature 45°C; Thermo Scientific) at a maximum pressure of 8 x 107 Pa with a linear gradient of 4% to 45% acetonitrile in 0.1% formic acid for 65 minutes. One MS full scan (resolution of 60,000 for a m/z 390-12,10) was performed, followed by MS/MS scans (resolution of 15,000) for the 15 most abundant ion signals. Precursor ions with 2 m/z isolation width were fragmented at a normalized collision energy of 30 using higher-energy collisional-induced dissociation (HCD). The automatic gain controls for the full MS scan was set to 3e6 and 1e5 for MS/MS. The DDA data was analyzed in MaxQuant (1.6.10.43) against a database comprising of *Homo sapiens* (UniProt proteome identifier UP000005640), common contaminants from other species, and iRT peptides (Escher et al., 2012). For the search, tryptic digestion with maximum of two missed cleavage was allowed. Carbamidomethylation (C) was set to static modifications, while oxidation (M) was set to variable modifications.

### Cell lines

Human monocytic cell line Tamm–Horsfall protein 1 (THP-1) (TIB-202, male; American Type Culture Collection) was cultured in RPMI 1640 medium (Sigma-Aldrich) supplemented with 10% FBS (Life Technologies) and 2 mM GlutaMAX (Life Technologies) at 37°C in 5% CO_2_. The cell density was kept between 0.2-1.0 × 10^6^ cells/ml with viability over 95% (determined with Erythrosin B (Sigma-Aldrich)), and cells were harvested at 0.5 ×10^6^ cells/ml for the phagocytosis assay.

### Phagocytosis Assay

The phagocytosis assay was performed and analyzed using the PAN method as described (de Neergaard et al., 2019). Briefly, phagocytosis was performed with 100 000 THP-1 cells in a final volume of 150 μl with different multiplicity of prey (MOP) from 0-400 at 37°C for 30 min with gentle shaking conditions. The bacteria had, as previously described, been fluorescently double-stained and opsonized in either 5% sera or 0.1 mg/ml Xolair. For the assessment across the different isolates, phagocytosis was performed at MOP 80, and the concentration of Xolair and IVIG was 0.5 mg/ml. Phagocytosis was halted by transferring samples to ice and kept cold during data acquisition using CytoFlex. At least 5 000 events of the population of interest were acquired. Ice samples were used as a control for internalization. Free bacteria were analyzed separately to determine the fluorescent intensity of a single bacterial unit and to confirm pH sensitivity of the staining pH was decreased by adding 1 μl of sodium acetate (3 M, pH 5.0). Four different colonies per isolate were assessed.

### Analysis of flow cytometry data

Flow cytometry data were analyzed using FlowJo version 10.6.2 (TreeStar). THP-1 cells were gated on forward (FSC) and side scatter (SSC) height. Events with extreme negative fluorescence were excluded. THP-1 cells positive for Oregon Green (FITC-H) were defined as associating, and those also positive for CypHer5E (APC-H) were defined as internalizing cells as well. Free bacteria were gated on SSC-H in combination with a positive Oregon Green signal, and doublets were excluded by gating on FSC-H versus FSC-A. Gating strategy is visualized in Supp. Fig 2.

The data was then further analyzed using the PAN method (de Neergaard et al., 2019) in Prism 9.3.1 (GraphPad Software). Inbuilt non-linear analysis tool “Agonist vs. response –Variable slope (four parameters)” was used to create curves and determine MOP_50_, corresponding to the MOP, which evoked half of the maximal response. One curve was generated per replicate, represented as a mean value in the figures. The persistent association was defined as the percentage of THP-1 cells positive for at least one bacterium either adhered or internalized. In comparison, internalization was defined as THP-1 cells positive for at least one internalized bacterium. Normalization for association was performed by interpolating the median fluorescence intensity (MFI) at MOP_50_.

For assessing the individual phagocyte ability, the amount of bacteria associated and internalized was determined by the MFI of an associating THP-1 cell. To convert it into the number of prey per phagocyte, PxP, each fluorescent signal (associated: Oregon Green, internalized: CypHer5E) was divided with the MFI of free bacteria for CypHer5E at pH 5. Since streptococci are typically not present as a single bacterium, a prey unit most likely represents a chain.

### Statistics

To compare the acute and convalescent serum effect on binding and phagocytosis of the corresponding isolate, the paired non-parametric Wilcoxon matched-pairs signed-rank test was used. No paring was performed if it had been normalized against acute sera, and the hypothetical value was set to 100. Two-way ANOVA with Šídák’s multiple comparisons test was applied when the sera were tested across the different isolates. The alpha value was set to 0.05. For the statistical tests Prism, version 9.3.1 (GraphPad Prism) was used, while data collection and simple calculations were performed using Microsoft Excel 2021 (Microsoft Corporation).

## Results

### Clinical characterizations of patients and bacterial isolates

Four patients (patients A-D (PA-PD)) with *S. pyogenes* bacteremia were enrolled. Their clinical characteristics are presented in Figure 1A. In summary, two females (PB, PC) were infected with isolates of *emm1,* and two males with *emm118* (PA) or *emm85* (PD), respectively. The primary infection foci were skin and soft tissue. Upon admission, all patients were hemodynamically stable, but within 48 h, patient D acquired septic shock (SEPSIS-3 criteria, (Singer et al., 2016)). Patient D’s symptoms commenced two weeks before admission, in contrast to the others that almost immediately were admitted to the hospital, suggesting that the serum from this patient could be in a different immune response phase compared to the other acute samples. The patients’ immunoglobulin (IgA, IgG, IgM) concentration in serum changed between admission (acute) and six weeks later (convalescent), with the largest difference being an increase in IgG for patients C and D (PA: 9 %, PB: 13 %, PC: 87%, PD 55 %) (Fig. 1B). To determine the subclass distribution of the immunoglobulins, we performed a mass spectrometric analysis of the patients’ sera (Fig. 1D). After infection, IgG1 increased for three out of four patients, whereas patient D had more IgG1 in the acute sera. Additionally, for patient C, the IgG distribution shifts to a noticeably higher level of IgG3 in convalescent sera. The *S. pyogenes* isolates were sequenced (Supp. Data), and the M protein sequences were compared (Fig. 1C). The M1 isolates (*S. pyogenes* from patient B, SpB, and patient C, SpC) had identical M proteins, but the overall similarity between M1, M85, and M118 was relatively low (30%). However, there was a higher sequence similarity between M85 and M118 when pairwise aligned (59.3 %). In summary, our patient group was small, with different ages and disease severity but similar health status. The infecting agents consist of three different *emm*-types, with two patients infected by the same type.

**Figure 1.**
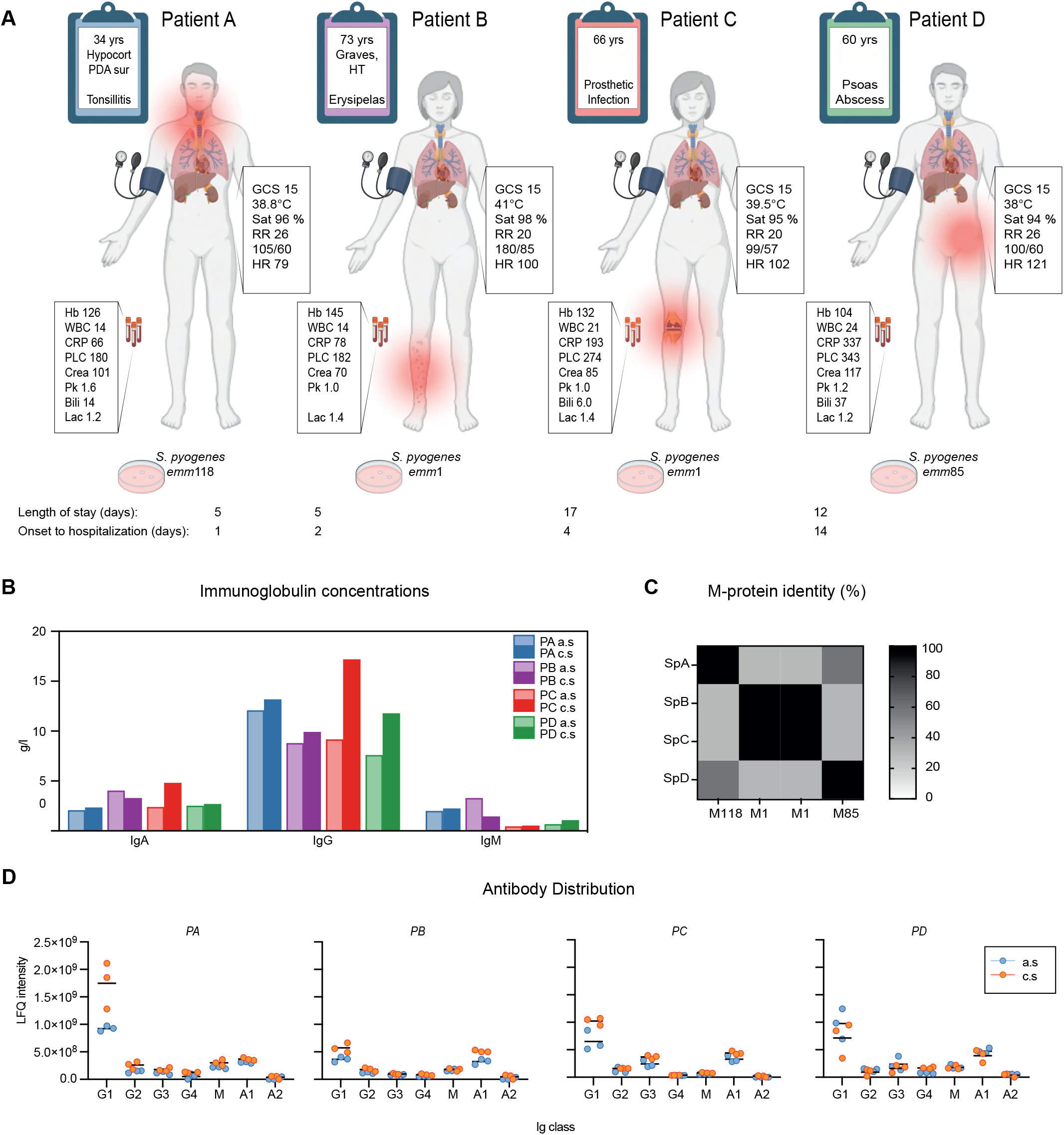
The characteristics of the patients and their infecting isolate. (A) The four patients (PA-PD) with invasive *S. pyogenes* infection included in the study. GCS, Glasgow Coma Scale; RR, respiratory rate; Sat, saturation; blood pressure mmHg; HR, heart rate; Hb, hemoglobin g/L; WBC, white blood count x10^9^/L; CRP, C-reactive protein mg/L; PLC, platelet count x10^9^/L; Crea, creatinine μmol/L; PK(INR), prothrombin complex international normalized ratio; Lac, lactate mmol/L. (B) The immunoglobulin concentration of acute (a.s) and convalescent (c.s) sera as reported from the clinical diagnostics data. (C) A multiple sequence alignment and identity percentage of the four detected M protein sequences, including two identical M1 sequences of SpB and SpC, M85-SpD, and M118-SpA.(D) The immunoglobulin (Ig) distribution was determined by mass spectrometry for each patient’s serum. The line represents the mean, n = 3. Figure A created with BioRender.com

### Antibody binding is increased after invasive *S. pyogenes* infection

To determine the effect an invasive *S. pyogenes* infection has on antibody binding, we opsonized each *S. pyogenes* isolate bacteria with the corresponding paired sera. The non-specific monoclonal antibody Xolair was a control for Fc binding, and IVIG was a positive control. To evaluate the contribution of the M protein as an antigen, we included an M protein-deficient *S. pyogenes* mutant, SF370dM (dM). When analyzing the whole curve in the convalescent sera, all patients had a significant increase in IgG bound to the bacteria (Fig. 2A), and already at 0.1 % serum, the differences could be detected (Fig. 2C). Interestingly, the amounts of IgG bound to dM increased from low at low serum concentration to almost the same level as the clinical isolates when measured at the higher concentrations (Fig. 2B). The result thus indicates the presence of both IgGs with high affinities against M proteins and additional antigens in the absence of M protein. To evaluate the relative change of bound IgG’s within each patient’s sera, the level of IgG bound to the bacteria in convalescent serum was expressed as a fold change to the level of IgG’s bound in acute serum (Fig. 2D). The M1 patients (PB, PC) had the largest relative change among the isolates, while patients A and D had almost no difference. Nonetheless, the dM strain had the highest relative increase for each patient.

**Figure 2.**
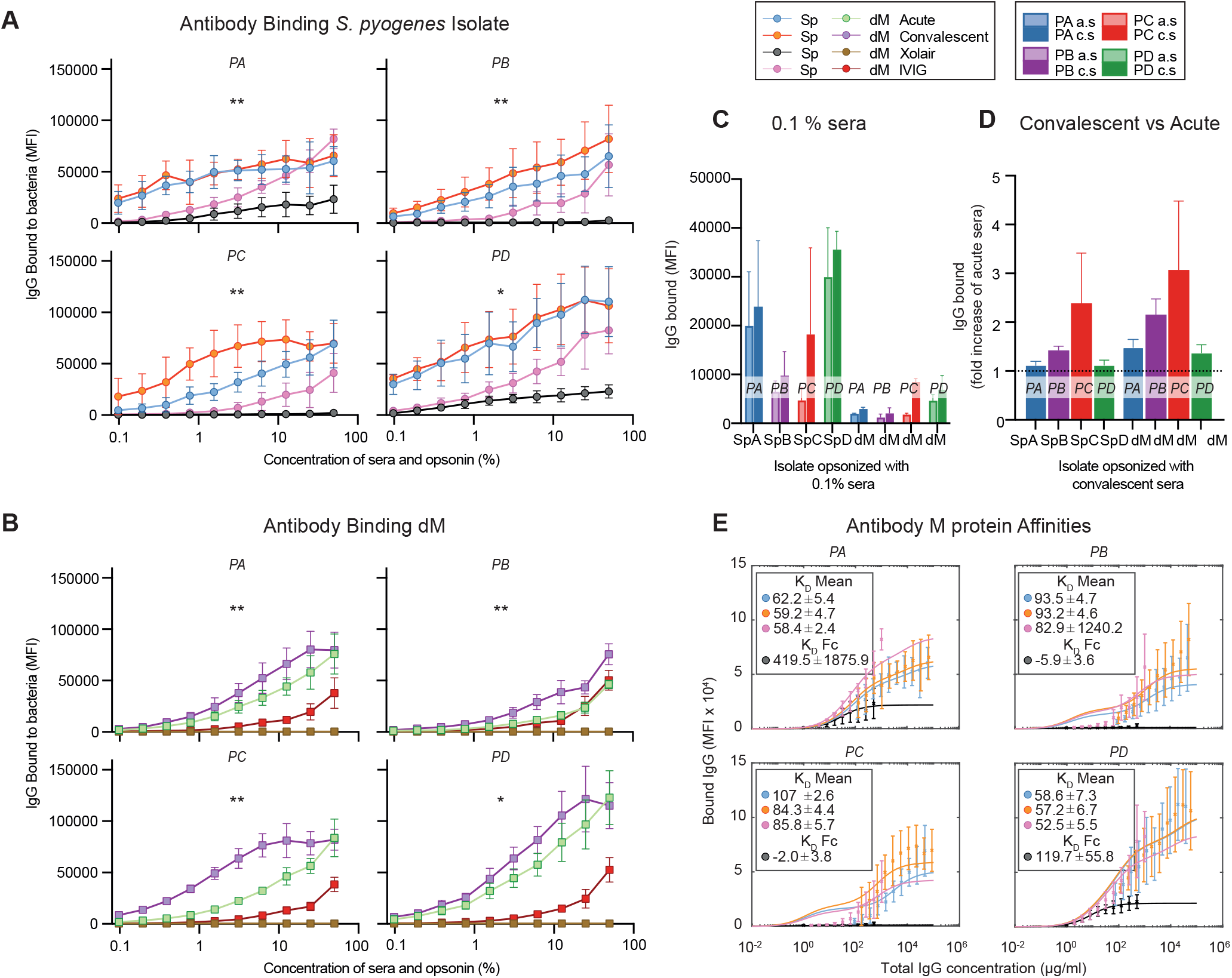
Antibody binding and distribution after invasive GAS infection. (A-E) The acute and convalescent sera from four patients (PA-PD) with *S. pyogenes* invasive infection were assessed. (A-D) The different isolates (Sp) and a strain lacking the M protein (dM) were opsonized in serial dilution of the sera and controls (see color legend). The undiluted concentration was 2 mg/ml for the polyclonal IgG (IVIG) and 1 mg/ml monoclonal non-specific IgG (Xolair). Binding was determined with IgG Fab-specific far-red fluorescent antibodies. Data were acquired through flow cytometry and are presented as mean ±SD, n = 4. (A-C) The amount of IgG bound to the bacteria isolate (A) or dM (B) for each opsonin expressed in fluorescent intensity of the secondary antibody. Wilcoxon matched-paired signed-rank test was performed on acute vs convalescent binding; p-value < 0.05 *, < 0.01 **. (D) The change in binding for convalescent compared to acute serum expressed in fold increase of bound IgG in acute serum. The baseline set at 1, marked with a line. © Affinities of each serum to M-protein through modeling of binding data. The affinity for sera and IVIG expressed in K_D_ mean log10 reference nM^-1^ while for Xolair, the M-protein Fc affinity is determined, named K_D_F_c_ with nM^-1^ as the unit.

To quantify the affinities of the binding IgG against the pathogen, we analyzed the binding curves using a bacteria-antibody binding model (Kumra Ahnlide et al., 2021) (Fig. 2E and Supp. Fig. 1A). Patient C had the highest increase in affinity after infection. However, both patients A and D had higher affinities than patient C already in acute sera. To summarize, antibody binding was increased to the infecting isolate after an invasive *S. pyogenes* infection regardless of *emm* type.

### Invasive infection leads to a differential opsonic response

To evaluate the dynamics of phagocytosis, we studied it from low to high bacteria-to-cell ratios (multiplicity of prey, MOP). By heat-inactivating the sera, we excluded contribution from the complement system to focus on Fc-mediated phagocytosis. We assessed the phagocytic ability of the phagocyte population based on the portion of phagocytes that can associate with or internalize their prey (defined as cells with at least one internalized bacterial unit) (Fig. 3A-B). In the convalescent serum, the association and internalization were significantly increased for patients A and C compared to acute sera. On average, 50% more phagocytes were associated with bacteria (Fig. 3A), and 75% more phagocytes had internalized at least one bacterium with patient C convalescent serum (Fig. 3B). The increase for patient A was 3% in association and 25% internalization. Patient D’s acute serum mediated higher association, whereas there was no difference in internalization as compared to convalescent serum. For patient B, there was no significant increase in neither association nor internalization. For dM, there was a significant increase from acute to convalescent sera for each patient in association ability and for all, except patient D, in the internalization ability. This indicates that the increase seen in phagocytosis when comparing convalescent to acute sera can not only be explained by antibodies binding to the M protein.

**Figure 3.**
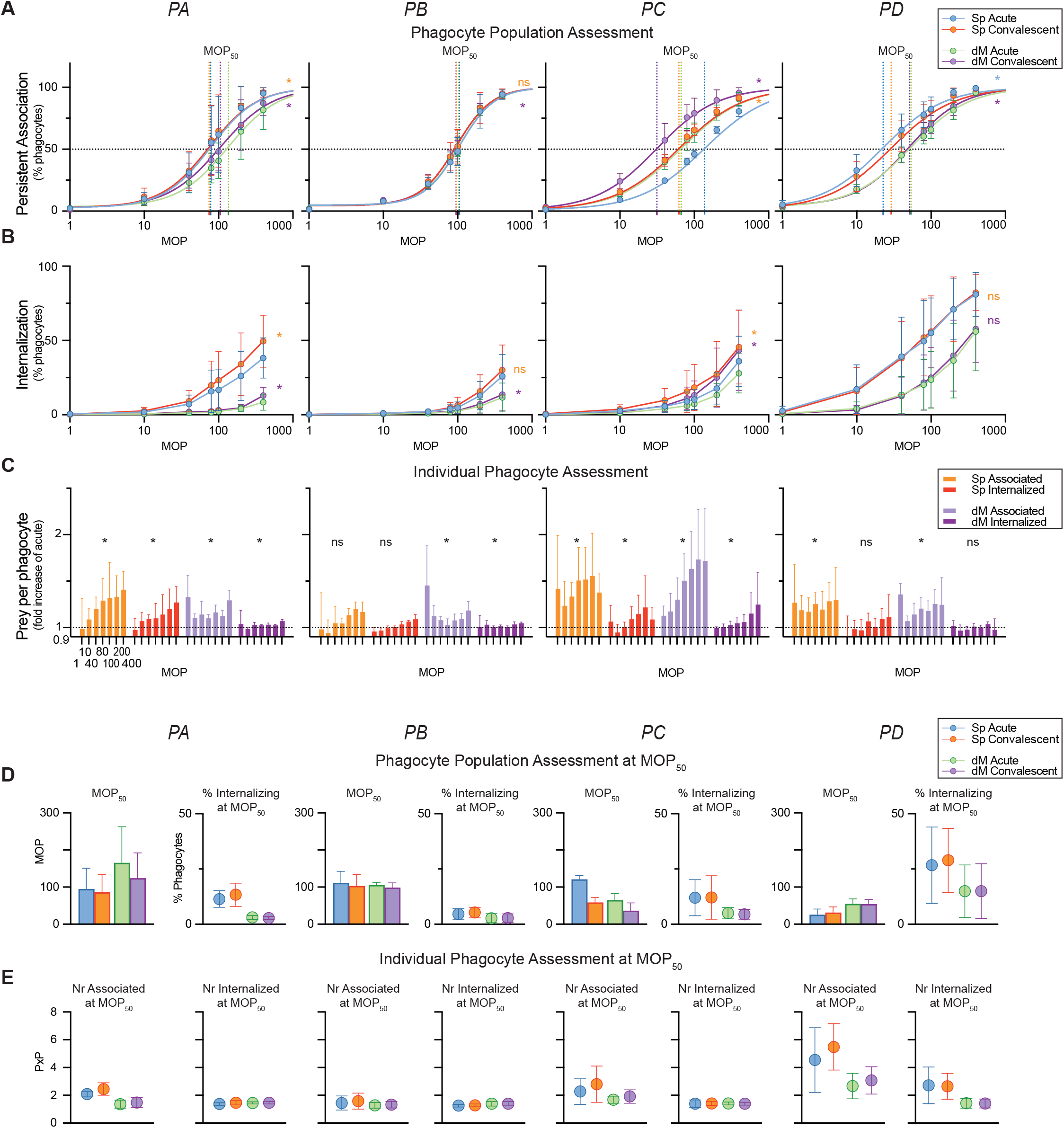
Assessment of the phagocytic response during infection. (A-E) The acute and convalescent sera from four patients (PA-PD) with *S. pyogenes* invasive infection were assessed. THP-1 cells were incubated (30 min, 150 μl, MOP 0–400, 37°C) with either the infecting isolate (Sp) or a strain lacking M-protein (dM) after the bacteria were fluorescently doubled stained with a pH-stable (Oregon Green) and a pH-sensitive (CypHer-5E) dye and opsonized in 5 % of the corresponding sera. Data were acquired through flow cytometry and are presented as mean ±SD, n = 4. (A-B) The percentage of phagocytes in the phagocytic population associating with (A) and internalizing (B) opsonized bacteria for each patient. In A, the average of the fitted persistent association curves is shown, and 50 % association is marked with a line for each curve. Acute vs. convalescent samples were compared with Wilcoxon matched-paired signed-rank test p-value < 0.05 *. (C) The change in the number of bacteria an individual associating phagocyte has associated with and internalized in convalescent compared to acute serum expressed as fold increase of acute serum. The baseline is visualized with a line at 1. The MOP range is 1-400. Wilcoxon signed-rank test was performed p-value < 0.05 *. (D) To the left, the MOP when 50 % of the phagocytic population have associated, called MOP_50_, based on the curves in A. To the right, the percentage of phagocytes internalizing at MOP_50_ is shown. (E) The average number of prey units per phagocyte, PxP, and associated with to the left and internalized to the right at MOP_50_.

In Figure 3C, by analyzing bacterial fluorescence intensities at a single-cell level, we provide an assessment of the phagocytic ability of each phagocyte, meaning its individual ability to adhere to and internalize bacteria. Compared to acute serum as the baseline, an increased association was detected for patients A, C, and D at all MOPs (p<0.05, median increase in % A: 29, C: 42, D: 25) but only at the highest concentration for patient B. Internalization, on the other hand, was only significantly improved for patient A and C (10% and 8.8%, respectively). For all the convalescence sera, dM was significantly more associated with cells than acute sera.

In Figure 3D, we demonstrate a comparison of the association capacity between the patients’ sera in a standardized manner by determining at what MOP 50% of the phagocyte population was associated with bacteria (MOP_50_). Patients A, B, and C require similar MOP, around 100, while patient D needs less than half to reach 50% association. Only patient C has a clear improvement in association capacity with convalescent sera (the MOP_50_ was halved). At MOP50, serum from patient D not only had the highest number of bacteria interacting with each phagocyte (Fig. 3E) but also the highest bacterial internalization in the phagocytes (Fig. 3D). When comparing convalescent to acute serum, patients A and D slightly increase the portion of phagocytes internalizing bacteria (Fig. 3E). For patients A, C, and D, the individual phagocyte adheres to more bacteria in convalescent sera, while internalization is unaffected. There are no differences for patient B and dM samples on the population or individual level.

When summarizing the different parameters analyzed, patient C had the most evident improvement in phagocytosis both on population and individual phagocyte level after infection; patient A had some improvement, while patient B had none. Patient D has the highest opsonic ability overall, which remains in the convalescent serum.

### Infection-induced opsonic antibodies are cross-reactive while non-opsonic response seems to be *emm* type-specific

To determine whether our findings were *emm* type-specific, we evaluated the effects of heat-inactivated sera on binding and phagocytosis across isolates. Overall trends are shown as heat maps (Fig. 4A, 4C), with quantitative analysis in Figures 4B and 4D. The IgG binding was significantly increased in the convalescent sera compared to acute sera across the different isolates, except for SpA (*S. pyogenes* isolate infecting patient A) opsonized with patient D sera where the increase was more modest (Fig. 4A-B). As expected, we see the highest binding to the infecting isolate for sera from patients A and D, but interestingly not for sera from patients B and C, which both were infected with *emm1* strains. In addition, the *emm1* isolates had fewer antibodies bound independent of which sera were tested. The *emm*1 isolates were also the least phagocytosed by each patient’s serum (Fig. 4C-D). However, there was a significant increase in phagocyte association for all the isolates opsonized in sera from patients A and C. In contrast, only SpA and SpD were improved for patient B, whereas patient D had no significant increase at all. Similar trends were seen with the antibody controls IVIG and Xolair, but with overall lower levels then the patient sera.

**Figure 4.**
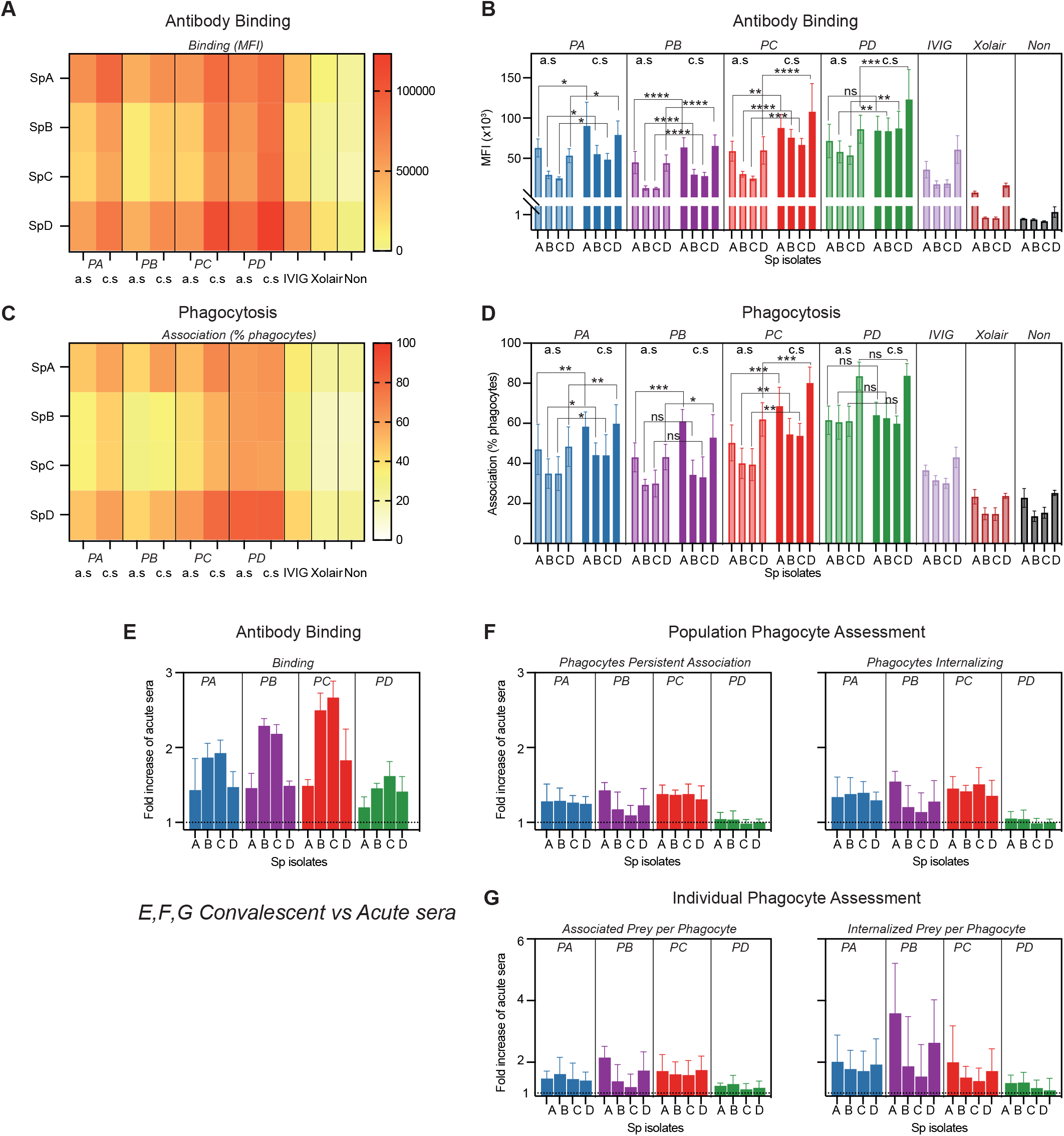
Antibody binding and opsonic function across strain for each patient. (A-G) The acute (a.s) and convalescent (c.s) sera from four patients (PA-PD) with *S. pyogenes* invasive infection were assessed. The different isolates (Sp) were fluorescently doubled stained with a pH stable (Oregon Green) and a pH-sensitive (CypHer-5E) dye and opsonized in 5 % of each serum. The concentration was 0.5 mg/ml for polyclonal IgG (IVIG) and monoclonal non-specific IgG (Xolair) and without opsonin for negative control. Binding was determined with IgG Fab-specific far-red fluorescent antibodies. Phagocytosis was performed with THP-1 cells incubated (30 min, 150 μl, MOP 80, 37°C) with each isolate. Data were acquired through flow cytometry and are presented as mean ±SD, n = 4. (A-D) Heatmaps and histograms visualizing IgG bound to each isolate (A-B) and the percentage of phagocytes associating with the bacteria (C-D). Significance tested with two-way ANOVA and Šídák’s multiple comparisons test, p-value < 0.05 *, <0.01 **, 0.001 ***, 0.0001 ****. (E-G) The change in binding (E) and phagocytosis (F-G) for convalescent compared to acute serum for each patient expressed as fold increase of corresponding acute serum. The baseline is visualized with a line at 1. (F) Visualizing the change in what portion of the phagocyte population associated and internalized bacteria while E looks at the change in the individual phagocytes capacity to associate (to the left) and internalize (to the right) bacteria.

In Figure 4 E-G we compare the convalescent sera relative to the acute sera. The largest increase in the antibody binding was against the *emm*1 isolates (SpB, SpC) for all patients, and patient C had the largest increase in antibody binding for all isolates (Fig 4E). The response in phagocytosis, both at the population (Fig. 4F) and the individual phagocyte level (Fig. 4G), was elevated in the same manner as the phagocyte association (Fig. 4C-D). Thus, after infection, patient B developed antibodies that were opsonic against isolates of other M types but non-opsonic against its infecting *emm*1 isolate (SpB). Nonetheless, serum from patient C, also infected with *emm*1, had increased binding and function against all isolates, including SpB, in convalescence. Hence, opsonic antibodies can be generated after an *emm*1 infection. Serum from patient A, infected by the *emm*118 isolate, increased antibody binding and phagocytosis across the strains, indicating a broad and improved opsonic antibody response after infection. Serum from the *emm*85-infected patient D maintained a high level of phagocytosis across strains, with no further increase, but with increased levels of antibody binding after infection (Fig. 4 E-G). To summarize (Table 1), after invasive *S. pyogenes* infection, cross-strain opsonic antibodies can be developed. On the other hand, non-opsonic binding antibodies are generated against specific types, and here it is primarily seen with the *emm*1 type.

**Table 1.**
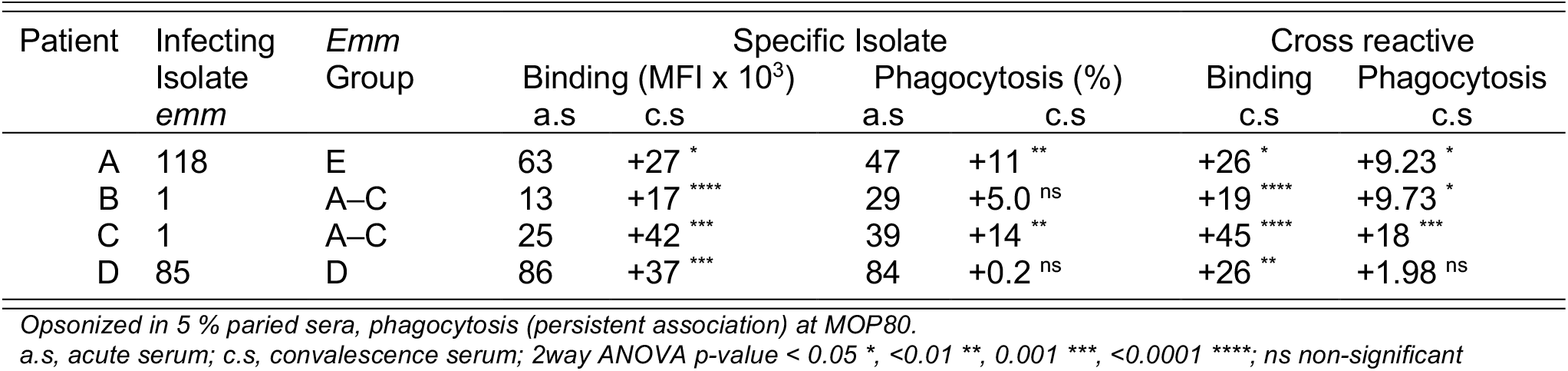
Adaptive response after *S.pyogenes* infection

## Discussion

The generation of opsonizing antibodies is a vital step in developing pathogen-specific immunity. In the present work, we have quantified the opsonic capacity of serum from patients during and after invasive *S. pyogenes* infection. Our results show that a modest increase in antibodies binding to the bacteria occurs after infection. It should be pointed out that this is from a relatively high basal level, as seen when compared to acute samples. However, this increase in binding antibodies does not always lead to an improved functional response in terms of phagocytosis. These findings are consistent with previous studies on *S. pyogenes* immunity, where antibodies binding to the conserved region of the M protein typically did not result in a bactericidal effect (Bahnan et al., 2021; Jones & Fischetti, 1988). One could speculate that the patient generating non-opsonic antibodies (PB) had an immune deficiency. Still, she generated opsonic antibodies toward other *S. pyogenes* types and had no disease record supporting that hypothesis. It was also not specific for *emm*1 type infection since the other patient infected with the *S. pyogenes* of the identical type did generate opsonic antibodies. The mechanism behind these two different responses is unknown, and it might be difficult to draw too firm conclusions from a limited set of patients. Still, we speculate that *S. pyogenes* might have mechanisms influencing the immune system so that it generates non-functional antibodies. Our results show that it is important to properly assay both antibody titers and antibody function to characterize an immune response.

Interestingly, the opsonic antibodies generated by the patients were cross-reactive and enhanced phagocytosis across types. Even if patients suffering from *S. pyogenes* invasive infection rarely get reinfected (Rasmussen, 2011), suggesting broader protection, *S. pyogenes* immunity is typically described to be type-specific (Jones & Fischetti, 1988). After an invasive infection, patients are expected to have a strong and specific antibody response rather than the modest and cross-reactive response seen in patients studied here. However, during the last decade, studies have reported the development of broadly opsonic antibodies in animal vaccine trials (Dale et al., 2011) and after superficial skin infections in school children (Frost et al., 2017). Furthermore, we recently found a protective human-derived antibody with opsonic function across a broad range of *emm* types (Bahnan et al., 2021). Here, we have provided clinical data on the development of cross-type opsonic antibodies after invasive *S. pyogenes* infection, which to our knowledge, has not previously been described.

Nevertheless, generalizations based on our results should be made with care since this study is based on a small study population. Still, the different infecting types in this study (emm1, emm85, emm118) belong to three diverse *emm* groups, A–C, D, and E, respectively (McMillan et al., 2013), so it is reasonable to describe the opsonic response in our patients as broad. Taken together with already published data, we believe this study provides further proof of the development of a general rather than a strictly type-specific *S. pyogenes* immunity after infection.

## Acknowledgments

This study acknowledges the Department of Clinical Microbiology, Office for Medical Services, Region Skåne, Lund, Sweden. We also thank Sebastian Wrighton and Arman Izadi for their important technical contribution. TdN was funded by the Royal Physiographic Society. PN was funded by the Swedish Research Council (VR).

---

**Supplementary Figure 1.** (A) Quantifying the affinities of each serum to M-protein through modeling of binding data. The affinity for sera and IVIG expressed in K_D_ mean log10 reference nM^-1^ while for Xolair, the M-protein Fc affinity is determined, named K_D_F_c_ with nM^-1^ as the unit.

**Supplementary Figure 2.** (A-B) Gating strategy of flow cytometry data for phagocytes (A) and bacteria (B) using FlowJo. (C-D) The clinical *S. pyogenes* isolates (Sp) opsonized with 0.5 mg/ml IVIG with phagocytosis (C: association, D: internalization) either on ice or 37°C. Data were acquired through flow cytometry and are presented as mean ±SD, n = 4.

